# P2RX7 inhibition reduces breast cancer induced osteolytic lesions - implications for bone metastasis

**DOI:** 10.1101/2021.12.31.474644

**Authors:** Karan M. Shah, Luke Tattersall, Aleana Hussain, Sarah C. Macfarlane, Alexander Williamson, Adelina E. Acosta-Martin, Janine T. Erler, Penelope D. Ottewell, Alison Gartland

**Affiliations:** The Mellanby Centre for Musculoskeletal Research, Department of Oncology and Metabolism, The University of Sheffield, Sheffield S10 2RX, UK; Biomolecular Science Research Centre, Department of Bioscience and Chemistry, Sheffield Hallam University, Sheffield S1 1WB, UK; biOMICS Facility, Faculty of Science Mass Spectrometry Centre, The University of Sheffield, Sheffield S10 2TN, UK; Biotech Research and Innovation Centre (BRIC), University of Copenhagen (UCPH), 2200 Copenhagen, Denmark

**Author notes:** Corresponding author: Prof. A. Gartland; Post: The Mellanby Centre for Musculoskeletal Research, Department of Oncology and Metabolism, The University of Sheffield, Beech Hill Rd, Sheffield, S10 2RX, United Kingdom.

**Keywords:** Bone, Breast cancer, Extracellular Vesicles, Hypoxia, Metastasis, P2RX7, Premetastatic niche, Purinergic signalling

## Abstract

Breast cancer metastasis to bone is a major contributor to morbidity and mortality in patients and remains an unmet clinical need. Purinergic signalling via the P2X7 receptor (P2RX7) in the primary tumour microenvironment is associated with progression of several cancers. It has also now become evident that intra-tumoural hypoxia facilitates cancer metastasis and reduces patient survival. In this study, we present data suggesting that hypoxia regulates the expression of P2RX7 in the primary tumour microenvironment; and importantly, inhibition with a selective antagonist (10mg/kg A740003) increased cancer cell death via apoptosis in a E0771/C57BL-6J syngeneic murine model. Furthermore, micro-computed tomography demonstrated reduced number of osteolytic lesions and lesion area following P2RX7 inhibition in absence of overt metastases by decreasing osteoclast numbers. We also demonstrate that activation of P2RX7 plays a role in the secretion of extracellular vesicles (EVs) from breast cancer cells. Mass-spectrometric analyses showed a distinct protein signature for EVs derived from hypoxic compared with normoxic cancer cells which elicit specific responses in bone cells that are associated with pre-metastatic niche formation. Thus, inhibiting P2RX7 provides a novel opportunity to preferentially target the hypoxic breast cancer cells preventing tumour progression and subsequent metastasis to bone

## 1. Introduction

Breast cancer is the most common cancer affecting females, with a worldwide incidence of nearly 2 million and is responsible for >600,000 deaths every year. Metastasis to distant organs account for 90% of these cancer-related deaths, with bone, lung and liver as the most common sites for metastases [1,2]. Metastatic bone disease is a major contributor to morbidity and mortality in cancer patients, with 70-80% of late stage breast cancer patients developing metastases in this site and five-year survival rate, following diagnosis of bone involvement dropping to just 13% (11 to 14) [3].

Patients with bone metastases suffer from skeletal complications such as bone pain, impaired mobility, pathological fractures, spinal cord compression and hypercalcaemia that lowers their quality of life [4-7], and this is a significant financial burden to the healthcare services [8]. Bone metastases remain incurable, with current treatment options being palliative, and slowing the progression of cancer. Whilst bone metastases occurs predominantly in hormone-receptor positive breast cancer patients, nearly 13% of patients with initial bone metastasis are hormone-receptor negative and 8% of these have triple-negative breast cancer [9]. Understanding mechanisms that prepare the bone microenvironment for colonisation by metastatic cells is vital for the development of therapeutics to prevent metastatic bone disease and improve the clinical outcome for these patients.

Intra-tumoural hypoxia is increasingly being recognised as an important regulator of tumour progression. It is a negative indicator for patient prognosis, and recent studies have shown that activation of hypoxic pathways are associated with poor survival in breast cancer [10-12]. Hypoxic signalling in breast cancer has been shown to increase tumour angiogenesis [13], facilitate epithelial-to-mesenchymal (EMT) transition [14,15], and increase tumour progression and metastasis [16-18]. The Lysyl oxidase (LOX) family of copper-dependant amine oxidases are consistently overexpressed in hypoxic tumours and are associated with poorer clinical outcomes [19-22]. We have previously demonstrated LOX expression is associated with increased bone relapse in breast cancer and facilitates the formation of pre-metastatic niche (PMN) in bone [19].

Hypoxia has also been implicated in regulating the expression of the purinergic P2X7 receptor (P2RX7) in breast cancer [23]. P2RX7 is an ATP-gated ion channel, which at low agonist concentrations is permeable to K^+^, Na^+^ and Ca^2+^ ions; but with sustained activation by higher extracellular ATP concentrations, forms a non-selective macropore permeable to large molecules up to 900Da [24]. The tumour microenvironment (TME) is characterised by high extracellular ATP concentrations and the expression of P2RX7 is upregulated in several cancers [25-27]. There is now increasing evidence of its role in cancer progression by supporting tumour growth, metastasis [28-30] and regulating cancer-associated inflammation [31].

In breast cancer, P2RX7 expression has been reported to be higher in the tumour tissue compared to the adjacent normal tissue [32,33]. P2RX7 has also been shown to stimulate invasion and migration of the oestrogen receptor positive T47D breast cancer cells via activation of the AKT pathway [34]. In a more recent study, P2RX7 has been implicated in promoting cell invasiveness by inducing changes in cell morphology via F-actin reorganisation, and formation of filopodia in 4T1 and MDA-MB-435 triple-negative breast cancer (TNBC) cells [35]. However, the direct role of P2RX7 in breast cancer metastasis to bone remains unexplored.

P2RX7 receptor activation is central to the assembly and induction of NLRP3 inflammasome and mediates the maturation and secretion of pro-inflammatory cytokines, including interleukin-1β (IL-1β) which has recently been shown to play a role in breast cancer metastasis to bone [36-39]. P2RX7 activation has also been shown to facilitate the secretion of extracellular vesicles (EVs) from breast cancer cells [40]. With LOX and LOX-like proteins having been shown to be secreted in EVs [41-43], P2RX7 activation in the primary tumour may have implications for EV-mediated PMN formation and distant metastasis to bone.

Here, we present novel data that suggests targeting P2RX7 in breast cancer may reduce the formation of pre-metastatic osteolytic lesions. This effect may partly be due to the reduced secretion of EVs from cancer cells that can alter bone homeostasis.

## 2. Results

### 2.1 P2RX7 is associated with poorer clinical outcomes in breast cancer

Analysis of RNA-seq expression data from The Cancer Genome Atlas (TCGA) and the SCAN-B datasets (N=4669) [44,45] revealed that P2RX7 expression is higher in patients with the aggressive basal-like breast cancer compared to the other molecular subtypes [46] (P<0.05; Figure 1A). Expression of P2RX7 was also significantly higher in triple negative breast cancer (TNBC) compared to non-TNBC patients (P<0.0001; Figure 1B) and this was associated with reduced distant-metastasis free survival when comparing tercile 1 and 3 (HR=1.59; P = 0.036; Figure 1C)[47].

**Figure 1.**
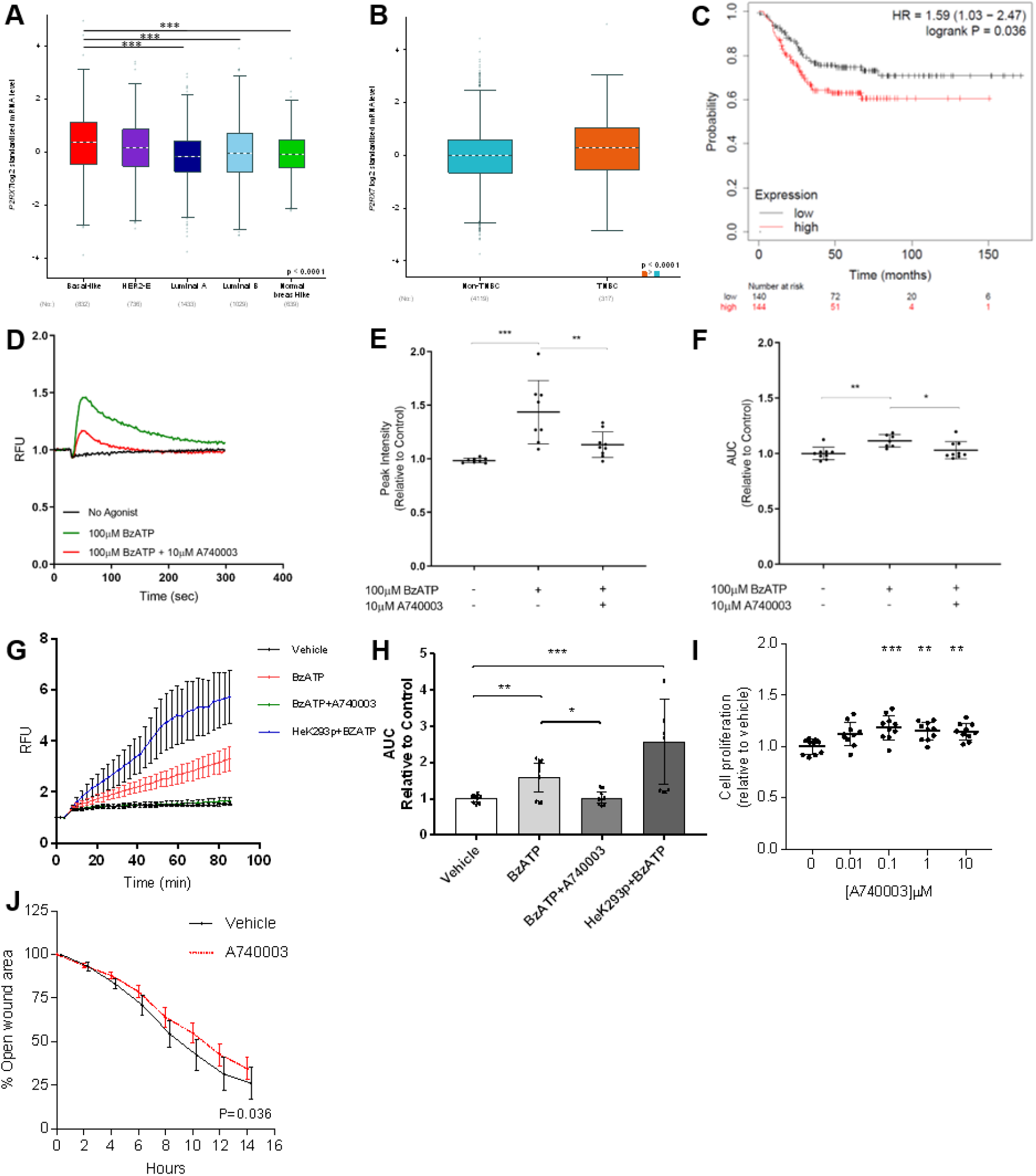
A) mRNA expression for P2RX7 in the different molecular subtypes of breast cancer. Data analysed from the TCGA and SCAN-B datasets (n=4669). B) mRNA expression of P2RX7 in triple-negative breast cancer (TNBC, n=317) compared to non-TNBC (n=4119). C) Kaplan-Meier plot for distant-metastasis free survival for patients with low (tercile 1) and high (tercile 3) P2RX7 expression. D) P2RX7 agonist BzATP (100μM) mediated calcium response (green) in E0771 murine breast cancer cells, which is inhibited in presence of a specific antagonist A740003 (10μM, red). Peak intensity and the area-under-curve (AUC) for the calcium response in (D) are presented in (E) and (F) respectively. G) P2RX7 mediated pore formation with higher concentrations of BzATP (300μM) assessed by ethidium bromide uptake in E0771 cells (red), which is inhibited by A740003 (green). P2RX7 over-expressing HEK-293 cells are used as a positive control (blue). (H) Analysis of the AUC for these pore-formation responses. I) Effect of P2RX7 antagonist A740003 on E0771 cell proliferation assessed by WST-1 at 72h. J) The effect of A740003 (10μM) on E0771 migration using scratch assay. All *in vitro* data is presented relative to the untreated controls and are from 3 separate biological repeats. The data has been presented as mean ± SD and analysed using one-way ANOVA. *P<0.05, **P<0.01 and ***P<0.0001.

### 2.2 Murine E0771 breast cancer cells express functional P2RX7 which alters cell proliferation and migration *in vitro*

Expression of functional P2RX7 was determined in the murine E0771 triple-negative breast cancer cells [48] by measuring the intracellular Ca^2+^ response (for ion channel function) and ethidium bromide uptake in cells (for pore-forming function) following agonist stimulation. Treatment with the specific P2RX7 agonist (100μM BzATP) caused a robust intracellular Ca^2+^ response compared to vehicle controls, which was inhibited by 10μM A740003 – a P2RX7 antagonist (Figure 1D). The peak intensity for the BzATP induced Ca^2+^ response (mean±SD; 1.43±0.29 vs 1.0±0.02; P<0.01; Figure 1E) and the area under curve (AUC) (1.11±0.05 vs 1.0±0.05) were reduced following antagonist treatment (1.13± 0.11 and 1.02±0.07 respectively; P<0.05; Figure 1F). Stimulation with higher concentrations of BzATP (300μM) resulted in P2RX7 mediated pore formation in E0771 cells, with HEK-293 cells transfected with P2RX7 serving as a positive control (Figure 1G). The AUC for agonist treated E0771 cells was larger than the vehicle controls (1.58±0.53 vs 1.0±0.11; P<0.01; Figure 1H) and was inhibited with A740003 antagonist (1.03±0.19; P<0.05; Figure 1H) demonstrating a P2RX7 mediated effect. Under normoxic conditions P2RX7 antagonist (A740003) at concentrations of (0.1-10μM) increased cell proliferation of E0771 cells after 72h treatment (P<0.05; Figure 1I). *In vitro* migration of the E0771 cells, as measured by a scratch assay, was reduced following antagonist treatment compared to the vehicle controls (P<0.05; Figure 1J).

### 2.3 Effects of P2RX7 inhibition on the primary tumour in a syngeneic model of breast cancer

Having observed an effect of P2RX7 on cell proliferation and migration and an upregulation in its expression under hypoxia, we next investigated the effects of P2RX7 inhibition in an immune-competent, metastatic E0771-C57BL/6J syngeneic model. The tumour burden assessed by measuring the bioluminescence signal from luciferase expressing E0771 cells showed no difference between the vehicle and A740003 (10mg/kg) treated mice (3.42×10^8^±3.84×10^8^ photons/sec/cm^2^/steradian vs 5.46×10^8^±6.45×10^7^; P>0.05; Figure 3A). There were no differences observed in the tumour volume between the two groups measured at the end of experiment (0.49±0.45cm^3^ vs 0.57±0.19; P>0.05; Figure 3A).

**Figure 2.**
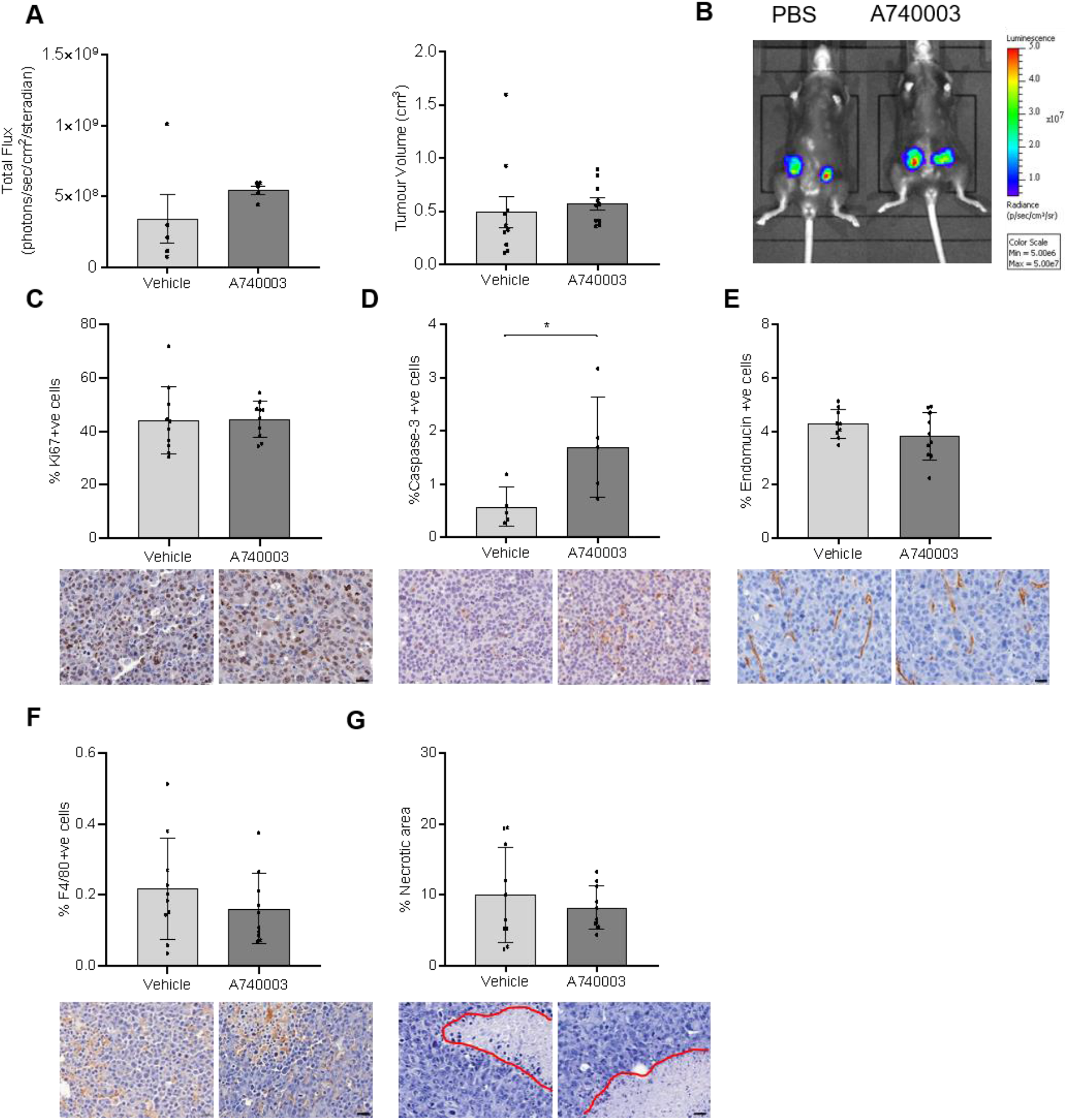
The effect of P2RX7 inhibition on the primary tumour. C57BL/6J mice were injected with luciferase expressing E0771 cells intra-ductally and were treated intraperitoneal with 10mg/kg A740003 or vehicle controls, daily (n=5 per group). The primary tumour growth was measured using A) bioluminescence–based *in vivo* imaging and Vernier callipers. B) Representative bioluminescence images of the primary tumours. At the end of the study, the effects of the A740003 treatment was assessed on the excised primary tumours by IHC. The percentage of C) Ki67 positive proliferating cells, D) caspase-3 positive apoptotic cells, E) endomucin positive endothelial cells, and F) F4/80 positive macrophages were measured by QuPath software. G) The percentage of necrotic area in the primary tumours (red outline) was assessed as an average of 3 levels, 100μm apart. The data has been presented as mean ± SD and analysed using unpaired t-test. *P<0.05. Scale bar = 20μm.

**Figure 3.**
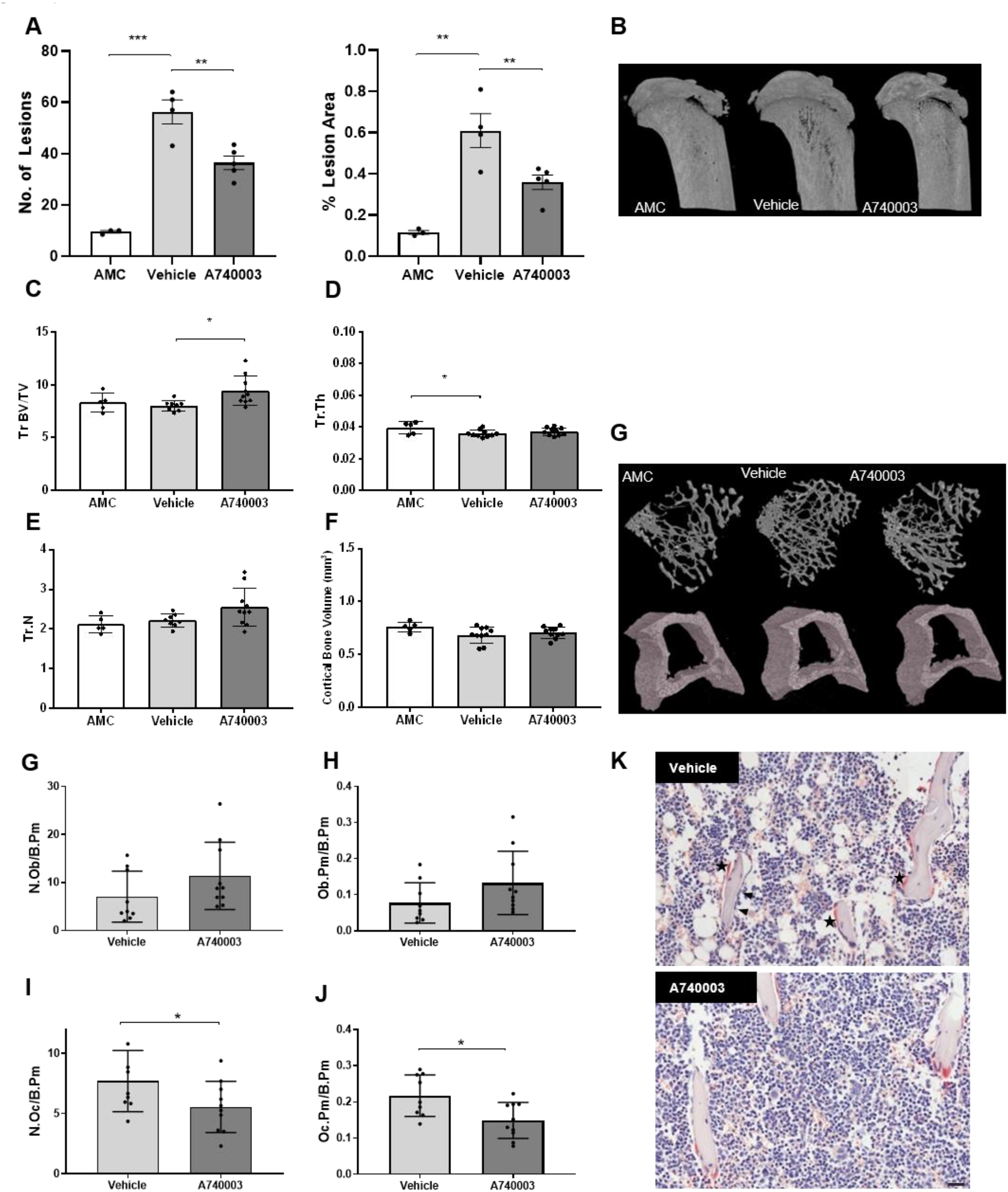
The effect of P2RX7 inhibition on the bone microenvironment. A) μCT analyses of osteolytic lesions in tibias of E0771 tumour bearing C57BL/6J mice treated daily with P2RX7 antagonist (10mg/kg A740003) or vehicle controls (n=5 per group); AMC are non-tumour bearing age-matched controls. B) Representative μCT images of proximal tibias from each group. Morphometric analyses of the proximal tibia was done to characterise the structural changes in bone by measuring C) trabecular bone volume fraction (Tr BV/TV), D) trabecular thickness (Tr.Th), E) trabecular number (Tr.N), and F) cortical bone volume. G) Representative 3D μCT images of the tibial trabecular and cortical bone are shown alongside. Historphometric analyses of the tibial trabecular bone was performed by assessing G) osteoblast number per bone perimeter (N.Ob/B.Pm), H) osteoblast surface per bone perimeter (Ob.Pm/B.Pm), I) osteoclast number per bone perimeter (N.Oc/B.Pm), and J) osteoclast surface per bone perimeter (Oc.Pm/B.Pm). K) Representative images of the tibial sections with TRAP-positive osteoclasts (stars) and osteoblasts (arrows) shown on trabecular bone. Scale Bar = 50μm. The data has been presented as mean ± SD and analysed using unpaired t-test. *P<0.05, **P<0.01 and ***P<0.0001.

We then investigated the effects of P2RX7 inhibition on the primary tumour TME and observed no differences in percentage of Ki67 positive proliferating cells (44.13±12.7 vs 44.37±6.79; Figure 3C). However, the caspase-3 positive apoptotic cells were higher in the A740003 treated group compared to the vehicle controls (1.69±0.95 vs 0.56±0.36, P<0.05; Figure 3D). Tumour vascularisation assessed by measuring the percentage of endomucin positive endothelial cells was also unchanged between the two groups (4.28±0.54 vs 3.82±0.89; Figure 3E). There was no change in the F4/80 positive macrophage population within the primary tumour between the treatment and the vehicle control groups (0.21±0.14 vs 0.16±0.09; Figure 3F), and percentage area of necrosis also remained unchanged between the two groups (10.03±6.69 vs 8.23±3.05; Figure 3G).

### 2.4 P2RX7 inhibition reduces pre-metastatic lesions in bone

None of the mice in either the treatment or vehicle control group developed overt bone metastases, detectable by bioluminescence. However, the tibiae of tumour-injected mice from the vehicle control group had increased number of osteolytic lesions with a mean diameter threshold of 50μm (9.5±0.86 vs 56.25±9.28; P<0.0001; Figure 3A), and percentage lesion area (0.11±0.01 vs 0.60±0.16; P<0.01; Figure 3A) compared to non-tumour bearing age-matched controls (AMC). P2RX7 antagonist treated mice had 35% fewer osteolytic lesions (36.4±5.87; P<0.01), and a 41% decrease in percentage lesion area (0.35±0.07; P<0.01) compared to the vehicle controls (Figure 3A). Standard bone morphometry analysis of the tibia demonstrated that the mice treated with A740003 had 17% higher trabecular bone content (Tr BV/TV; 9.43±1.37) compared to the vehicle controls (8.03±0.50; P<0.05; Figure 3C); other parameters including trabecular thickness (Tr.Th), trabecular number (Tr.N) and cortical bone volume remain unchanged between the two groups (Figure 3 D-F). Histomorphometric analyses was performed by measuring osteoblasts and tartrate-resistant acid phosphatase (TRAP) positive osteoclasts on the endocortical and trabecular bone surfaces of the tibia. Whilst there were no changes in the bone cell populations on endocortical surfaces between the two groups (Supplementary Figure 1A-B), A740003 treated mice had 28% lower osteoclast number per millimetre of trabecular bone (N.Oc/B.Pm; 7.71±2.53 vs 5.56±2.13; P<0.05; Figure 3I) and 33% less bone surface occupied by osteoclasts (Oc.Pm/B.Pm; 0.21±0.05 vs 0.14±0.04; P<0.05; Figure 3J).

To investigate if the effects of A740003 observed in this study were due to the direct effects of P2RX7 inhibition on bone cells, we performed experiments with mature primary murine osteoclasts (post-onset of resorption) and human osteoblast-like SaOS-2 cells *in vitro*. There were no changes observed in total number of osteoclasts, number of resorbing osteoclasts, percentage area of resorption and the resorption ability of osteoclasts following treatment with A740003 (Supplementary Figure 1C-G). Exposure to A740003 did not affect the percentage area of mineralisation by SaOS-2 cells *in vitro* (Supplementary Figure 1H).

Lower number of osteolytic lesions and percentage lesion area were observed following A740003 treatment in a 4T1-BALB/C syngeneic model, with mice that lacked P2RX7 (Supplementary Figure 2A). This further suggests that the effects of A740003 were at least partly due to P2RX7 inhibition in the primary tumour.

### 2.5 Activation of P2RX7 in E0771 cells mediates EV secretion that alter bone homeostasis

EVs secreted by cancer cells have been shown to mediate organotropism and PMN formation [49]. To determine if the observed effects of P2RX7 inhibition on bone parameters in the absence of overt metastasis were mediated by cancer derived EVs, we investigated the effects of hypoxia and P2RX7 inhibition on EV secretion. We isolated EVs from E0771 cells using size-exclusion columns (fractions 5-8; Figure 4A), and nanoparticle–particle tracking confirmed the median particle size to be 102 nm (Figure 4B). Immunoblotting for markers of EVs (CD9 and TSG101) further confirmed the isolated fractions to be enriched in EVs, when compared to whole cell lysate and post-EV fractions (Figure 4C).

**Figure 4.**
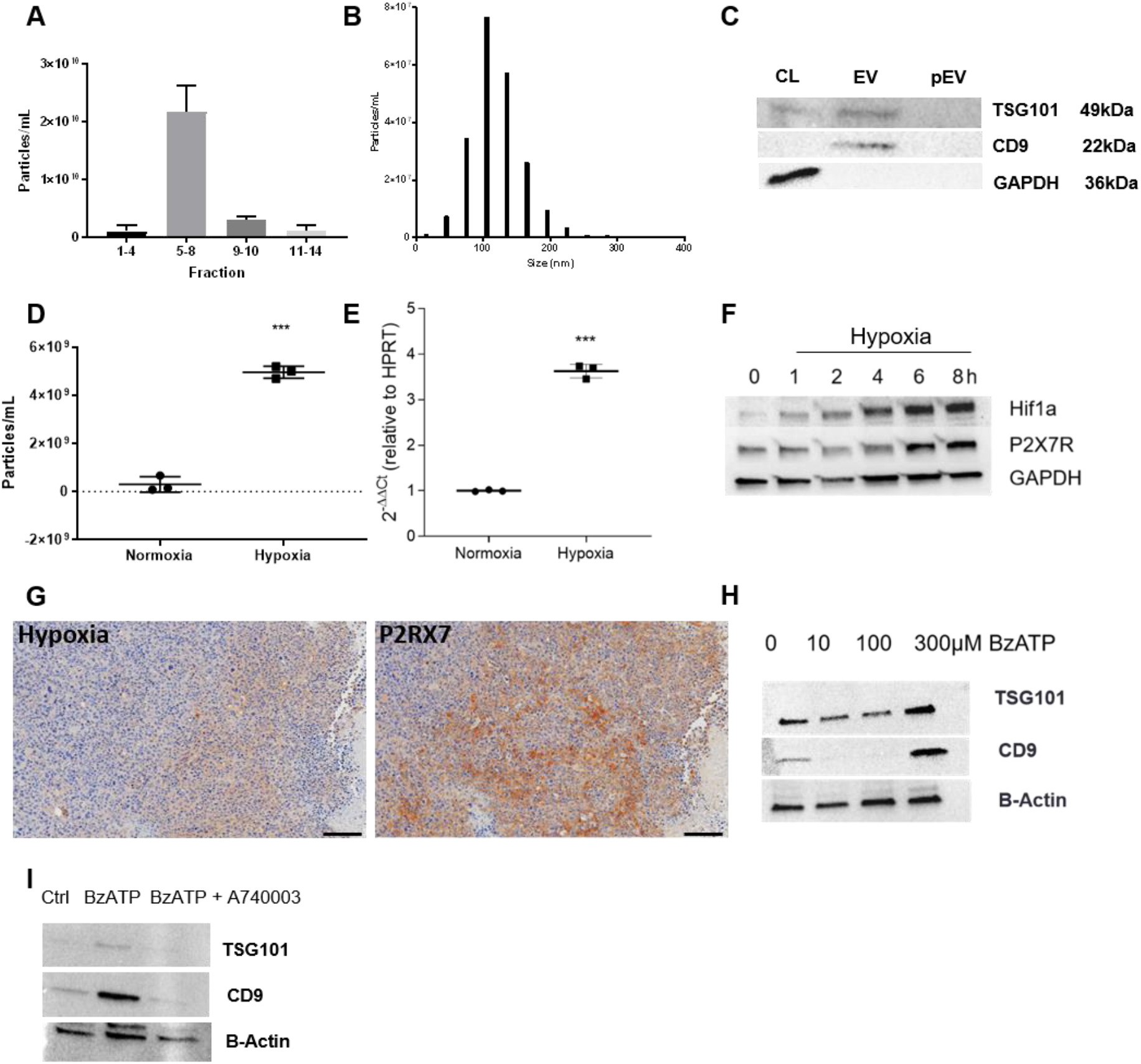
Extracellular vesicles (EVs) from E0771 cells were isolated using size-exclusion columns. A) EVs were enriched in elute fractions 5-8 which were pooled together for subsequent studies. B) size distribution of the isolated EVs as measured by nanoparticle tracking analyses (NTA). C) Western blotting for CD9 and TSG101 as markers for EVs in the cell lysates (CL), EV fractions and post-EV (pEV) fractions. E0771 cells were cultured in hypoxia (1% O_2_) or normoxia for 8h and EVs isolated subsequently from the conditioned media. D) NTA of the isolated EVs from hypoxic and normoxic cells. E) P2RX7 expression in E0771 cells cultured in normoxic and hypoxic conditions for 8h as assessed by real-time PCR. F) Hif1a and P2RX7 protein expression over time under hypoxic conditions with GAPDH as a loading control. G) The effect of hypoxia on P2RX7 expression was also assessed in primary tumours *in vivo*. IHC staining for GLUT-1 (as a marker for hypoxia) correlates with P2RX7 in serial sections. H) E0771 cells were treated with varying concentrations of BzATP (0-300μM), and EVs isolated from the media. Western blotting for CD9 and TSG101 shows the effect of P2RX7 activation on EV secretion, with β-Actin serving as a loading control. I) A740003 mediated inhibition of P2RX7 activation by 300μM reduces the EVs (CD9 and TSG101) secreted in the conditioned media. All *in vitro* data are from 3 separate biological repeats. The data has been presented as mean ± SD and analysed using unpaired t-test. ***P<0.0001.

Following acute hypoxia (1%O_2_ for 8h), E0771 cells released 65% more EVs compared to cells grown in normoxic conditions (3.00×10^8^±3.2×10^7^ particles/mL vs 4.96×10^9^±3.5×10^7^; P<0.0001; Figure 4D). We next investigated if hypoxia regulates P2RX7 expression and show that acute hypoxia (1%O_2_ for 8h) led to a marked upregulation in P2RX7 transcript expression compared to normoxic E0771 cells (3.62±0.15 vs 1.00±0.01; P<0.0001; Figure 4E). This hypoxia mediated increase in transcript expression also translated to increased protein expression as measured by Western blotting (Figure 4F). This increased expression of P2RX7 *in vitro* was also seen *in vivo* as areas of hypoxia in our tumours models *in vivo* were associated with P2RX7 expression and GLUT-1 staining (being used as a marker for hypoxia [50] (Figure 4G)). Furthermore, P2RX7 activation by BzATP had a biphasic effect on EV secretion, with a decrease observed at lower concentrations (10 and 100μM), and an increase at 300μM (Figure 4H). This increase in EV release at the higher concentrations was reduced by inhibiting P2RX7 with 10μM A740003 (Figure 4I).

Additionally, E0771 cells stimulated with BzATP caused a dose-dependent increase in mature-LOX secretion in the conditioned media (Supplementary Figure 2B), and this increase was reduced following P2RX7 inhibition with A740003 (Supplementary Figure 2C).

To understand if these cancer-derived EVs could mediate the formation of osteolytic lesions, we assessed their effects on survival and function of bone cells *in vitro*. Primary murine osteoclasts on dentine disks exposed to hypoxic and normoxic EVs (100 particles/cell) did not show any changes in osteoclast number, number of resorbing osteoclasts, percentage area of resorption, or their resorption ability compared to vehicle control (Figure 5A-D). The viability of osteoblast-like SaOS-2 cells (Figure 5E) was also unaffected with exposure to hypoxic and normoxic EVs. However, the alkaline phosphatase (ALP) activity was lower following exposure to cancer-cell derived hypoxic (1.00±0.18) and normoxic EVs (1.08±0.20), compared to osteogenic media treated control conditions (1.42±0.27; P<0.0001; Figure 5F).

**Figure 5.**
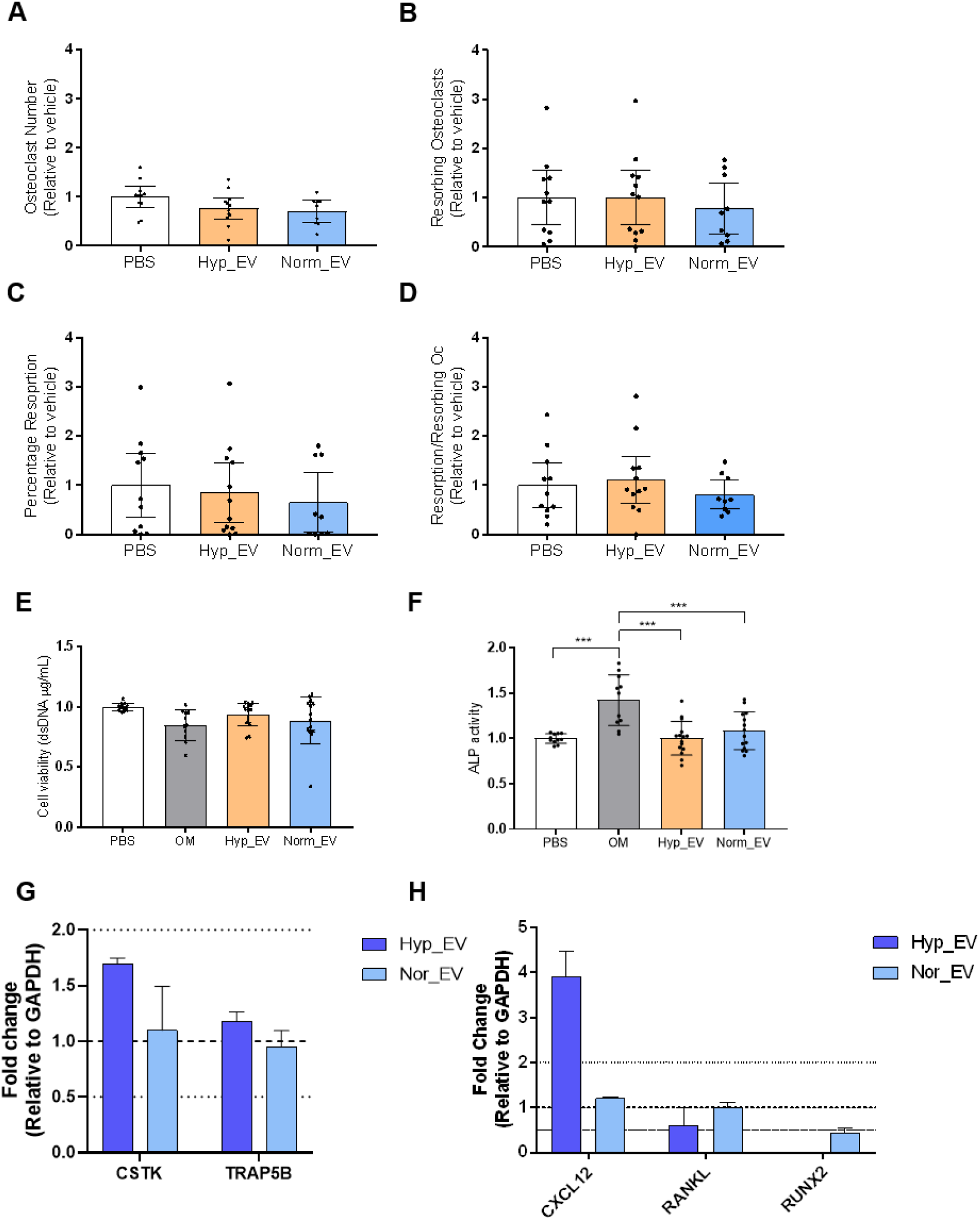
The effect of hypoxic and normoxic EVs on bone cells *in vitro*. Osteoclasts were generated from the mononuclear haematopoietic cell population isolated from long bone marrow of 8-week old female mice and cultured on dentine disks. The cells were treated with 100 hypoxic (Hyp_EV) or normoxic (Norm_EV) EV vesicles isolated from E0771 cells, per seeded cell for 14 days, with treatments replenished every 2-3 days. At the end of the culture, the cells were fixed, TRAP-stained and quantified for A) total osteoclast number, B) percentage resorbing osteoclasts, C) percentage resorption area, and D) resorption per resorbing osteoclast. SaOS-2 osteoblast-like cells were treated with 100 Hyp_EV or Norm_EV vesicles isolated from E0771 cells per seeded cell and E) viability, and F) alkaline phosphatase activity assessed at day 7. The effects of Hyp_EV and Norm_EV on expression of genes that affect G) osteoclast and H) osteoblast function following 24h exposure were assessed by RT-PCR. Log2FC ≥1 and ≤-1 (denoted by the horizontal lines) is considered to be significant. All data are from 3 separate biological repeats. The data has been presented as mean ± SD and analysed using One-wat ANOVA. ***P<0.0001.

Gene expression analysis for mature osteoclasts exposed to hypoxic and normoxic EVs showed that cathepsin-K (CTSK) and TRAP, which are associated with osteoclast function, were unchanged following exposure to two EV populations (Figure 5G), but distinctly different effects on gene expression was observed in osteoblast-like cells. Hypoxic EVs significantly reduced RANKL expression (FC=0.49) and increased CXCL12 expression (3.89) compared to controls (normalised to 1), whilst both normoxic and hypoxic EVs reduced RUNX2 expression (0.44 and 0.04 respectively) (Figure 5H).

### 2.6 Hypoxic EVs have a distinct proteomic expression profile that may mediate the observed effects on bone cells

To understand the differences in the protein cargo of EVs derived from hypoxic cells compared to those from normoxic cells, we performed MS analysis on the two EV populations. Hypoxic EVs had a distinct protein expression profile compared to the normoxic EVs and formed a separate cluster on the expression heatmap (Figure 6A). Furthermore, we observed that the hypoxic EVs contained 120 unique proteins that were not expressed in the normoxic EV population (Figure 6B listed in Supplementary Table 1). We next investigated if there was a difference in the expression of proteins present in both hypoxic and normoxic EVs. There were 56 proteins upregulated and 1 protein downregulated in the hypoxic EVs compared to the normoxic samples as illustrated in the volcano plot (Figure 6C; FDR=0.05; s0=0.1; Supplementary Table 2). A protein-protein interaction network was computed using STRING analysis for the upregulated proteins connections at a medium confidence level (Figure 6D). The analyses showed an upregulation in proteins enriched in the ‘ribosome’ (KEGG:mmu03010; FDR=2.6e-8), ‘viral carcinogenesis’ (KEGG:mmu05203; FDR=3.3e-2) and ‘PI3K-Akt signalling’ (KEGG:mmu04151; FDR=4.4e-2) pathways.

**Figure 6.**
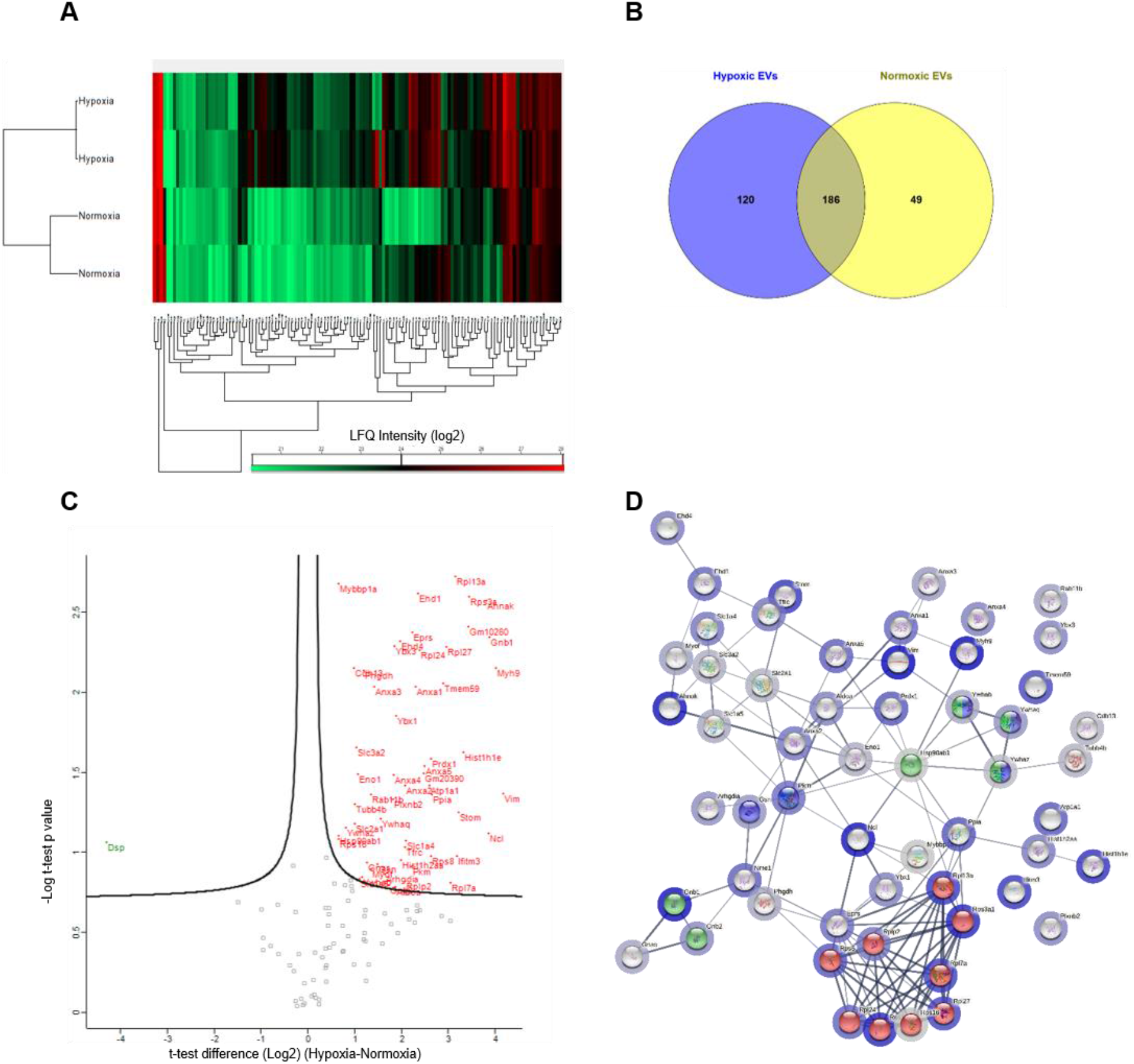
Proteomic analyses of hypoxic and normoxic EV sub-populations. A) Heatmap and hierarchical clustering for protein LFQ intensities in normoxic and hypoxic EVs. B) Venn diagram illustrating unique and common proteins present in normoxic and hypoxic EVs. C) Volcano plot for differentially expressed proteins. Proteins significantly upregulated in hypoxic EVs are shown in red, and downregulated are in green. D) The proteins upregulated in hypoxic EVs are analysed by STRING to show protein-protein connections at medium confidence. The halo around the node represents the fold change in protein expression compared to normoxic EVs with darker blue corresponding to higher values. The thickness of the edges correspond to the confidence level of the interactions. The data presented here has been analysed from 2 separate biological experiments.

## 3. Discussion

Breast cancer metastasis to bone remains incurable and is associated with significant patient morbidity and mortality. In this study, we present novel data that suggests a role of P2RX7 in facilitating bone metastasis. Specifically, we show that P2RX7 is upregulated in breast cancer under hypoxic conditions, and its inhibition reduces osteolytic lesions in the absence of overt metastasis by reducing osteoclast activity. Furthermore, P2RX7 activation regulated the secretion of EVs from cancer cells, and EVs secreted from the hypoxic cancer cells altered gene expression in osteoblasts and thus may deregulate bone homeostasis.

To validate the clinical relevance of P2RX7 in breast cancer, we analysed publicly available clinical datasets and show that *P2RX7* has higher expression in the more aggressive basal-like and triple-negative breast cancer compared to luminal or normal breast-like subtypes. This increased expression was also associated with poorer distant metastasis free-survival. Whilst clinical evidence for the role of *P2RX7* in patient prognosis for breast cancer is limited, its increased expression is associated with lower survival in non-small cell lung cancer [51], gastric cancer [52], neuroblastoma [53], and clear-cell renal cell carcinoma [54]. *In vitro* and *in vivo* studies have implicated P2RX7 in breast cancer cell invasion and metastasis from both ER+ve and TNBC subtypes [34,35], and this is consistent with the lower distant metastasis free survival associated with high *P2RX7* expression in patient population observed here.

In this study we used the E0771 murine breast cancer cells from the C57BL/6 background, which forms spontaneous metastasis in a syngeneic model. We confirmed the presence of functional P2RX7 by assessing both ion channel activity and pore forming ability using BzATP as a specific agonist, and A740003 a potent selective and competitive antagonist, at previously reported concentrations [55,56]. P2RX7 inhibition by A740003 increased E0771 cell proliferation, consistent with a recent study by Brisson *et al*. (2020) that showed P2RX7 knock-out 4T1 breast cancer cells had increased proliferation under basal conditions, at levels similar to those observed here [35]. Also consistent with previous reports on the effects of P2RX7 on cancer cell invasion and migration [34,40], we observed reduced cell motility for 10μM A740003 treated cells. Taken together, these data further support a role for P2RX7 in facilitating breast cancer progression.

Hypoxia in the tumour microenvironment is being recognised as an indicator for poor patient prognosis and is associated with increased metastasis [10,12]. With P2RX7 activation implicated to confer a more metastatic phenotype in cancer cells, we investigated if the receptor expression is regulated by hypoxia. In 1% O_2_ conditions, we observed a robust increase in P2RX7 expression at both mRNA and protein levels, which associated with increased HIF-1α. Furthermore, this increase of P2RX7 also translated to primary tumours *in vivo*, with areas of hypoxia associated with higher P2RX7 expression in serial sections. These findings are in agreement with the studies by Tafani et al. (2011), who demonstrated P2RX7 to be downstream of HIF-1α and to facilitate cell invasion via Nf-κB translocation in MDA-MB-231 and MCF-7 breast cancer cell lines [23].

The tumour-host interactions in the TME plays a vital role in tumour growth and progression and given the pivotal role of P2RX7 in modulating angiogenesis and immune responses [53,57,58], the *in vivo* effects of its inhibition are likely to be multifactorial. In our study, P2RX7 inhibition did not affect primary tumour burden and volume, but increased caspase-3 positive apoptotic cells. P2RX7 receptor is known to play a role in energy metabolism and has been shown to aid in cellular adaptations to adverse growth conditions including serum starvation and low glucose via metabolic reprogramming [59,60]. Furthermore, this P2RX7 mediated metabolic reprogramming has also been associated with increased cancer stemness, chemoresistance and ultimately metastasis [61,62]. It is therefore likely that P2RX7 confers a pro-survival adaptation under nutrient deprived hypoxic conditions that exist within the tumour mass, and inhibition of the receptor leads to increased apoptosis.

Pre-clinical evidence for the role of P2RX7 in breast cancer is currently limited. P2RX7 antagonists have shown a tendency to reduce tumour growth following the establishment of the primary tumour in a syngeneic model, with no effect on metastatic colonisation [35]. In a contrary report, P2RX7 inhibition was shown to reduce spleen and liver metastasis in tamoxifen-resistant MCF7 xenograft model [40]. Most reports describing the effect of P2RX7 inhibition in breast cancer are predominantly evidenced by *in vitro* studies and show an effect on cell migration and invasion [35,63].

Indeed, the lower number of osteolytic lesions in the P2RX7 antagonist treated group in absence of overt metastasis detectable by bioluminescence suggests a reduction in the metastatic cascade. This reduction in osteolytic lesions was associated with increased trabecular bone volume and reduced osteoclast number. The P2RX7 antagonist directly did not alter the mature osteoblast and osteoclast function *in vitro*. In addition, a similar reduction in osteolytic lesions was also observed with P2RX7 inhibition in a separate *in vivo* syngeneic model with 4T1 breast cancer cells injected in BALB/C host lacking P2RX7. Taken together, this suggests that the effect of the antagonist, is at least partly due to the inhibition of P2RX7 in the primary tumour. Given the absence of effect on tumour proliferation or vascularisation, it is possible that the P2RX7 inhibition reduces tumour cell migration and invasion, and ultimately the formation of micro metastasis that can deregulate the osteoblast-osteoclast coupling. The threshold for *in vivo* bioluminescent signal has been shown to be around 400 cells [64], and smaller metastasis could cause the observed effects. Alternatively, P2RX7 activation has been shown to facilitate release of various molecules and cytokines, and its inhibition may alter the secretion profile of the primary tumour thereby affecting the phenomenon of pre-metastatic niche formation in bone.

Recent studies have shown that EVs derived from the primary tumour mediate organotropism and facilitate the formation of pre-metastatic niches in target organs. Previous studies by us and others have shown that hypoxic secretome of the primary tumour facilitates the formation of PMN. Indeed, in this study we show that hypoxic cancer cells secrete more EVs than normoxic cells. Furthermore, the increase in EV secretion following activation of hypoxia– regulated P2RX7 at agonist concentrations that mediate pore formation, suggests a possible underlying mechanism for this effect. This is consistent with recent research suggesting a role of P2RX7 activation in the release of EVs in pathophysiological conditions [65]. P2RX7 activation also led to increased secretion of LOX in the conditioned media, a known mediator of PMN formation in bone. Therefore, targeting P2RX7 presents a novel opportunity to specifically target the hypoxic cancer secretome thus preventing PMN formation and delaying metastasis.

Although both hypoxia and P2RX7 have been associated with EV secretion previously [40], it was unclear if the hypoxic sub-population of EVs have a differential effect on the PMN formation in bone. We show that at physiologically relevant concentrations of EVs (100 particles/cell) osteoblast differentiation is reduced by both the hypoxic and normoxic EVs. Interestingly, the hypoxic EVs had a differential effect on osteoblast expression for *RANKL* and *RUNX2*, which have implications for bone homeostasis. Moreover, *CXCL12*, a chemokine associated with increased colonisation by cancer cells was markedly upregulated following hypoxic EV exposure [66]. CXCL12 or stromal-cell derived factor-1 (SDF-1) is expressed in the osteoblast/endosteal, the haematopoietic stem cell and the vascular endothelial niches, which are integral to the bone metastatic niche. Indeed, interactions between C-X-C chemokine receptor type 4 (CXCR4)/CXCL12 is thought to be a key modulator in the homing and adhesion of cancer cells to these metastatic niches within bone [67]. CXCR4 expressed on cancer cells mediates the attachment to CXCL12 (a homeostatic chemokine produced by the osteoblasts and the vasculature in bone), therefore the upregulation of CXCL12 observed here may have implications for tumour cell colonisation of bone and overt metastases [68-70].

Proteomic analysis confirmed a distinct expression profile for the two EV sub-populations and provide a basis to mechanistically understand these differential effects. Understanding the hypoxic EV protein signature can inform the development of prognostic biomarkers for more aggressive metastatic cancers and identify molecules that may mediate organotropic metastasis. For instance desmoplakin (Dsp), an integral component of desmosome structures that maintain cell-cell adhesion, was the only protein markedly downregulated in the hypoxic EVs. Downregulation of Dsp has already been suggested as a potential prognostic marker for breast cancer [71]. Conversely, non-muscle myosin heavy chain IIA (MYH9) upregulated in hypoxic EVs has been shown to promote cell invasion and metastasis in gastric cancer [72], drug resistance in neuroblastoma [73] and promotes a stem-cell like phenotype in lung cancer [74]. Similarly, vimentin (vim) a major constituent of the intermediate filament protein family was upregulated in hypoxic EVs and has been shown to correlate with well-established poor prognostic factors in breast cancer patient samples [75,76]. Protein-protein interaction analyses identified proteins from the ribosome and the PI3K-Akt pathways to be enriched in the hypoxic EVs. Whilst specific proteins such as RPL27a (ribosome pathway) and GNB1-2 (PI3K-Akt signalling) which were upregulated here, have been shown to play a role in development and metastasis of breast cancer previously [77,78], a more thorough investigation is warranted to understand their clinical significance and exploit the molecular mechanisms that may be involved.

Taken together our data suggests that P2RX7 expression is higher in hypoxic breast cancer cells and may give them a survival advantage. We have also shown that the activation of P2RX7 in these cells facilitates the secretion of pro-metastatic factors, including EVs and LOX, that may promote PMN formation in distant organs. Thus, inhibiting P2RX7 provides a novel opportunity to preferentially target the hypoxic breast cancer cells that facilitate tumour progression and distant metastasis.

## 4. Materials and Methods

### 4.1 Cell lines

Murine breast cancer cells E0771 (purchased from ECACC) and 4T1 (ATCC) were cultured in DMEM© GlutaMax™ media supplemented with 10% FBS and 1% Penicillin/streptomycin (hereon referred to as complete medium) (Gibco, Invitrogen, Paisley, UK) and maintained in a humidified incubator at 37°C and 5% CO_2_. Human osteoblast-like SaOS-2 cells were kindly provided by Professor Jim Gallagher (Liverpool, UK), and were also cultured in complete medium. All cell lines were authenticated by short tandem repeat profiling and regularly checked for mycoplasma contamination.

### 4.2 Fluo-4 Direct Calcium Assay

Changes in intracellular calcium response were measured using Fluo-4 Direct™ Calcium Assay (ThermoFisher Scientific, Paisley, UK) in accordance with the manufacturer’s instructions. Briefly, E0771 cells were seeded in a 96-well plate at a density of 15×10^3^ cells per well in complete media and left overnight to adhere prior to the assay. Cells were washed with PBS and were incubated with Fluo-4 Direct™ Calcium reagent for 1h at 37°C, with or without 10μM A740003 P2RX7 antagonist (Bio-Techne Ltd, Abingdon, UK). The cells were then stimulated with 100μM 2’(3’)-O-(4-Benzoylbenzoyl) adenosine-5’-triphosphate (BzATP; Merck Life Sciences, Gillingham, UK) to activate the P2RX7, and the fluorescence measured using FlexStation-3 Multimode Microplate reader (Molecular Devices, Wokingham, UK) with excitation at 494nm and emission at 506nm. Subsequently, the cells were treated with 500nM ionomycin to obtain the maximal fluorescence for normalisation of the data. The data was acquired for 300 seconds, including an initial 20 second baseline reading

### 4.3 P2RX7 mediated pore formation

Cellular uptake of Ethidium Bromide (EtBr) was used to assess the P2RX7 mediated pore formation in E0771 cells. Briefly, E0771 cells were seeded in a 96-well plate at a density of 15×10^3^ cells per well in complete media and left overnight to adhere prior to the assay. Cells were washed with PBS and incubated with or without 10μM A740003 in Hank’s Balanced Salt Solution (HBSS) (ThermoFisher Scientific, Paisley, UK) for 1h at 37°C. Subsequently, cells were stimulated with 300μM BzATP in HBSS containing 100μM ethidium bromide (Invitrogen, Paisley, UK). The induction of pore formation was measured using FlexStation-3 Multimode Microplate reader with excitation at 360nm and emission at 580nm. The data was acquired for 45min, including an initial 5min baseline, and readings taken every 2min.

### 4.4 Cell proliferation

To assess cell proliferation, the E0771 cells were seeded at a density of 5000 cells per well in a 96-well plate in complete media. After 24, the cells were washed and treated with increasing concentrations of A740003 in media supplemented with 0.5%FBS. 72h post treatment, cell proliferation reagent WST-1 (Merck Life Science, Gillingham, UK) was added to the wells, incubated at 37°C for 2h and the absorbance read at 450nm using SpectraMax M5e Microplate Reader (Molecular Devices, San Jose, USA).

### 4.5 Cell Migration

Cellular migration was assessed using scratch assays for which 12-well plates were seeded with 200,000 cells in complete medium and left overnight to form a confluent monolayer. The medium was then changed to complete medium containing 5μg/mL mitomycin C and cells incubated for 2h at 37°C. A scratch was then made using a 10μL pipette tip in the centre of the well. The wells were washed and supplemented with vehicle or 10μM A740003 containing media supplemented with 0.5%FBS, and images taken every 2h to monitor wound closure. The images were analysed using the automated T-scratch software [79].

### 4.6 RNA Extraction and Quantitative Real-time PCR (qRT-PCR)

Cells were washed twice with ice-cold PBS and total RNA extracted using the RNeasy Extractions Kits (Qiagen, Hilden, Germany) as per manufacturer’s protocol and the quality checked using the Agilent 2200 TapeStation (Agilent Technologies, Santa Clara, CA, USA). Subsequently, 1ug of RNA was converted to cDNA using the High-Capacity RNA-to-cDNA reverse transcriptase kit (ThermoFisher Scientific, Paisley, UK) as per the manufacturer’s protocol. For qRT-PCR, relative expression of candidate genes was quantified by TaqMan™ assays (Applied Biosystems, ThermoFisher, Paisley, UK) in accordance to manufacturer’s instructions using a 7900HT Real-time PCR machine (Applied Biosystems, ThermoFisher, Paisley, UK). The TaqMan™ assays used were: Hs00175721_m1 (*P2RX7)*; Mm00484039_m1 (*CTSK)*; Mm00475698_m1 (*ACP5*/*TRAP5B)*; Hs00171022_m1 (*CXCL12);* Hs00243522_m1 (*TNFS11/RANKL)*; Hs01047975_m1 (*RUNX2)*; Mm99999915_g1 and Hs04420632_g1 (*GAPDH*) (ThermoFisher Scientific, Paisley, UK).

### 4.7 *In vivo* studies

All animal studies were performed using 7-9 week old female C57BL/6J (Charles River, Margate, UK), or BALB/c P2RX7^−/-^ mice (kindly provided by Professor Niklas Rye Jørgensen, Department of Clinical Biochemistry, Rigshospitalet, Glostrup, Denmark) that were acclimatised for one week prior to experimental manipulation. All mice were housed in the same environmentally controlled conditions at 22°C, with a 12h light/dark cycle and were fed *ad libitum* with 2018 Teklad Global 18% Protein Rodent Diet (Harlan UK Ltd., Huntingdon, UK) and water. All procedures complied with the Animals (Scientific Procedures) Act 1986, and were approved by the local Research Ethics Committee at The University of Sheffield (Sheffield, UK). For the E0771-C57BL/6J syngeneic model, the mice were anesthetized by inhalation of isofluorane and 1.25 × 10^5^ luciferase expressing E0771 cells were engrafted via intra-ductal injection and immediately followed by injection of 0.003 mg Vetergesic. After 24h, the mice were randomised to treatment groups (N=5/group), and received either vehicle or 10mg/kg A740003 treatment injected intraperitoneal daily. The tumour growth was monitored by Vernier calliper measurements and bioluminescence–based *in vivo* imaging using IVIS Lumina II *In vivo* Imaging System and quantified by Living Image® software (version 4.5.4, PerkinElmer, Beaconsfield, UK) twice a week, 5min following subcutaneous injection of 6mg/kg D-Luciferin (PerkinElmer, Beaconsfield, UK). The experiment was ended when the tumour size reached ∼1cm^3^ in either the vehicle or the treatment group. At the end of the experiment, the mice were euthanised and the primary tumours and tibias collected and fixed in 10% neutral buffered formalin for *ex vivo* analyses. For the 4T1-BALB/c P2RX7^−/-^ studies, 0.5×10^5^ luciferase expressing 4T1 cells were engrafted via intra-ductal injection. After 24h, the mice were randomised to treatment groups (N=5/group), and received either vehicle or 10mg/kg A740003 treatment injected intraperitoneal daily. When the tumour size reached ∼0.5cm^3^, the mice were re-anesthetized and the primary tumours were surgically resected. The mice were followed a further 3 weeks, and at the end of the experiment, the mice were euthanised and tibias collected and fixed in 10% neutral buffered formalin for *ex vivo* analyses.

### 4.8 Immunohistochemistry and histological analyses

Immunohistochemical detection of GLUT-1 (Abcam, ab652), Ki67 (Abcam, ab15580), caspase-3 (R&D Systems, AF835), Endomucin (Santa Cruz, sc-53940), and F4/80 (Bio-Rad, MCA497R) were performed using the ABC system (Vector Laboratories, Burlingame, USA). The specific protocols used can be found in Supplementary Table 3. Briefly, after excision, tumours were fixed in 4% paraformaldehyde (PFA) overnight before embedding in paraffin according to standard histopathology techniques. Antigens were retrieved by incubating the slides in Citrate Buffer pH 6.0 at 95°C for 25min using a PT module (ThermoFisher Scientific, Paisley, UK). Endogenous peroxidase was then blocked by incubating slides in 3% H_2_O_2_ (Sigma-Aldrich, Gillingham, UK) in distilled H_2_O for 10min at room temperature. Slides were then blocked with 1.5% blocking serum (ABC VECTASTAIN IgG Kit) and incubated with the primary antibody (concentrations can be found in Table S1) at 4 °C overnight. After successive incubations (30min at RT) with the corresponding biotinylated IgG and the ABC solution (Vector Laboratories, Burlingame, USA), the peroxidase activity was developed using 3,3’-diaminobenzidine (DAB) (ImmPACT DAB EqV, Vector Laboratories, Burlingame, USA) as a substrate. Slides were counterstained with Gill’s hematoxylin (Merck Life Science, Gillingham, UK) for 20s, dehydrated, and mounted with coverslips using DPX (distyrene, plasticizer and xylene mixture) mounting medium (Sigma-Aldrich, Gillingham, UK). For assessing tumour necrosis, the tumour sections were stained with hematoxylin and eosin and percentage area of necrosis measured based on cell morphology. For each tumour sample, three sections 100μm apart were measured and an average percentage area of necrosis quantified. All stained slides were scanned using Pannoramic 250 Flash III slide scanner (3DHISTECH, Budapest, Hungary) and analysed using QuPath [80].

### 4.9 Micro-computed tomography of the bones

Formalin fixed tibiae were scanned using a SkyScan 1172 desktop micro-computed tomography (μCT) machine at a resolution of 4.3μm with the X-ray source operating at 50 kV, 200μA and using a 0.5 mm aluminium filter. Two-dimensional μCT images of the proximal tibiae were captured and reconstructed by Skyscan NRecon software at threshold of 0.0–0.14. The trabecular morphometry was characterized by measuring structural parameters in 1.0 mm thick region-of-interest, which was 0.2 mm below the growth plate. Cortical morphometry was quantified from the 1.0 mm thick cortical regions 1.0 mm below the growth plate. Nomenclature and symbols were used to describe the μCT derived bone morphometries according to the recommendations from Bouxsein et al. (2010) [81]. To quantitate cortical osteolytic lesions, a 3D model of the bone was constructed using Skyscan NRecon software (v. 1.6.9, Bruker, Belgium) and datasets were resized using Skyscan CTAn (v. 1.14.4, Bruker, Belgium). The datasets were imported into Osteolytica and lesion numbers and percentage lesion area analysed with a minimum lesion diameter of 50μm as threshold [82].

### 4.10 Bone Histomorphometry

For histomorphometrical analyses of the bone cells, the tibiae were fixed in 4%PFA solution, decalcified in 14.3% EDTA for 4 days at 37°C with daily changes of EDTA, then embedded in paraffin wax. Sections were cut (at 3μm) using a Leica Microsystems Microtome and stained with tartrate-resistant acid phosphatase (TRAP) as described previously [83]. The numbers of osteoblasts and TRAP-positive osteoclasts were determined on a 3mm length of endocortical surface starting 0.25mm from the growth plate and viewed on a DMRB microscope (Leica Microsystems). All histomorphometric parameters were based on the report of the American Society for Bone and Mineral Research histomorphometry nomenclature [84] and were obtained using the OsteoMeasure bone histomorphometry software (OsteoMetrics, Decatur, GA, USA). During preparation and analysis of tibiae, investigators were blinded to specific treatment groups.

### 4.11 Extracellular vesicle isolation

Size-exclusion chromatography (SEC) columns (IZON Science, Lyon, France) were used to isolate EVs from the cell culture conditioned media (CM). Briefly, 2×10^6^ cells were seeded in T75 flasks and cultured in complete medium for 2 days (80% confluence). The cells were then washed with PBS and phenol-free DMEM GlutaMax™ media supplemented with 0.5% exosome-depleted FBS (ThermoFisher Scientific, Paisley, UK) and 1% penicillin/streptomycin was added to the cells and incubated for 8h in normoxic (21%O_2_) or hypoxic (1%O_2_, maintained using H35 Hypoxystation, Don Whitley Scientific, Bingley, UK) environment. Subsequently, the CM was collected and centrifuged at 400g for 10min to remove any cell debris and the supernatant filtered through a 0.22μm filter. The CM was then centrifuged at 10000g for 10min and the supernatant concentrated using 10K MWCO protein concentrator columns (ThermoFisher Scientific, Paisley, UK). The concentrated CM was added to the qEV size exclusion columns (IZON Science, Lyon, France) and elute fractions enriched in EVs (5-8) collected as per the manufacturer’s protocol and pooled. The particle number and size distribution of the isolated EVs were determined by nanoparticle tracking analysis (NTA) using PMX-110 Zetaview® (Particle Metrix, Meersbusch, Germany) as per the manufacturer’s default software settings for EVs.

### 4.12 Western blotting

For EV samples, a total of 1μg of protein was used for the SDS-PAGE. The primary antibodies used were: TSG101 (1μg/mL, Abcam ab30871), CD9 (0.1μg/mL, Abcam ab92726), GAPDH (0.09μg/mL, Abcam ab181602), β-Actin (0.2μg/mL, Abcam ab16039), LOX (1:1000, Novus Biologicals NB100-2530H) and P2RX7 (3μg/mL, Alomone Labs APR-004). The membranes were blocked with 5% non-fat milk and 2.5% BSA for 2h and incubated with the primary antibody at 4°C overnight. The membranes were washed with TBS-T and incubated with corresponding HRP conjugated secondary antibody (Agilent Dako, Stockport, UK) for 1h at room temperature. The proteins were visualised by chemiluminescence using SuperSignal™ West Dura Substrate (ThermoFisher Scientific, Paisley, UK) on ChemiDoc XRS+ (Bio-Rad Laboratories Ltd., Watford, UK).

### 4.13 Osteoblast cell viability

SaOS-2 cells were seeded in a 96-well plate at a density of 5 × 10^3^ cells per well in 0.2 ml complete medium and left to adhere for the first 24h. The media was then replaced to DMEM© GlutaMAX™ containing 100U/ml penicillin, 100μg/ml streptomycin, 0.5% FBS, 10nM Dexamethasone and 50mg/mL L-Ascorbic acid (referred to hereon as osteogenic media) ± 5 × 10^5^ EV particles (hypoxic and normoxic) till day 7 at 37°C in a humidified atmosphere of 95% air and 5% CO_2_. The media and treatments were replenished every 2-3 days. Double stranded DNA was measured to assess cell viability, using the Quant-iT™ PicoGreen™ dsDNA assay kit (ThermoFisher Scientific, Paisley, UK) as per the manufacturer’s instructions.

### 4.14 Osteoblast Alkaline Phosphatase (ALP) Activity

SaOS-2 cells were seeded in a 96-well plate at a density of 5 × 10^3^ cells per well in 0.2 ml complete medium and left to adhere for the first 24h. The media was then replaced to DMEM© GlutaMAX™ containing 100U/ml penicillin, 100μg/ml streptomycin, 0.5% FBS, 10nM Dexamethasone and 50mg/mL L-Ascorbic acid (referred to as osteogenic media) ± 5 × 10^5^ EV particles (hypoxic and normoxic) till day 7 at 37°C in a humidified atmosphere of 95% air and 5% CO_2_. The media and treatments were replenished every 2-3 days. At day 7, the cells were washed with PBS and frozen with nuclease-free water at −80°C. ALP activity was measured by para-nitrophenyl phosphate (pNPP, Sigma–Aldrich, Dorset, UK) hydrolysis, and normalized to DNA content measured by Quant-iT™ PicoGreen™ dsDNA Assay Kit (ThermoFisher Scientific, Paisley, UK).

### 4.15 Bone Marrow Osteoclast Culture

Osteoclasts were derived from the mononuclear haematopoietic cell population isolated from long bone marrow of 8-week old female mice. The bone marrow from the limbs were flushed out by PBS using a syringe with 25-gauge needle. The cells were harvested by centrifugation and resuspended in selection medium of α MEM + GLUTAMAX™ (Gibco, Invitrogen, Paisley, UK), 100U/mL Penicillin and 100μg/mL Streptomycin, 10% FBS, and 30ng/ml M-CSF (R&D Systems, Abingdon, UK) and incubated in T75 flasks for 24h at 37°C, 5% CO_2_ to allow attachment of the stromal cells. Subsequently, the non-adherent cells were collected by centrifugation and resuspended in growth medium (α MEM + GLUTAMAX™ containing 100U/mL Penicillin and 100μg/mL Streptomycin, 10% FBS, 150ng/ml M-CSF, and 30ng/ml murine RANKL(R&D Systems, Abingdon, UK)) and were seeded onto dentine disks (generated in-house) in 96-well plates, at a density of 0.5 × 10^6^ cells per well and incubated overnight at 37°C, 7%CO_2_ to allow the attachment of osteoclast precursor cells. The wells were then washed once and cells were cultured in the growth medium ± 5 × 10^5^ EV particles (hypoxic or normoxic) for 14 days at 37 °C, 7% CO_2_ with the medium being replaced every 2– 3 days. At the end of the culture, the cells were fixed in ice cold 10% buffered formalin, TRAP stained and counterstained by Gill’s haematoxylin. The number of osteoclasts, resorbing osteoclasts (defined as a TRAP positive cell in or in close proximity to resorption pits) and the percentage area of resorption per dentine disk were quantified.

### 4.16 Proteomic sample preparation and liquid chromatography tandem mass spectrometry (LC-MS/MS)

S-trap (Protifi, Huntington NY, US) were used according to manufacture’s instructions to prepare 1ug of protein (as measured by bicinchoninic acid assay) of each EV sample. Disulphide bonds were reduced with 0.5M Tris(carboxyethyl)phosphine (TCEP) and alkylated using 0.5M iodoacetamide (IAA) prior to protein digestion with trypsin for 1hr at 47°C. Digested peptide samples were dried in a vacuum concentrator and resuspended in 40μl of 0.5% formic acid before LC-MS/MS analysis.

Peptides for each digested EV sample were separated in a Dionex Ultimate 3000 uHPLC (ThermoFisher Scientific, Paisley, UK) with an Acclaim™ PepMap™ 100 C18 trap column (3μm particle size, 75μmx150mm) and an EASY-Spray™ C18 column (2μm particle size, 50μmx150mm) using an 80min gradient at 0.25μl/min with 0.1% formic acid in water (mobile phase A) and 0.1% formic acid in 80% acetonitrile (mobile phase B) as follows: 0-5min, hold at 3% B; 5-60min, from 3% B to 40% B; 60-65min, from 40% B to 90% B; 65-70min, hold at 90% B; 70-71min, from 90% B to 3% B; 71-80min, re-equilibrate at 3% B.

MS/MS analysis was performed on a LTQ Orbitrap Elite (ThermoFisher Scientific, Paisley, UK) hybrid ion trap-orbitrap mass spectrometer equipped with an EASY-SprayTM Ion Source. MS survey scans in positive ion mode were acquired in the FT-orbitrap analyzer using an m/z window from 375 to 1600, a resolution of 60,000, and an automatic gain control target setting of 1×10^6^. The 20 most intense precursor ions were selected for the acquisition of MS/MS spectra in the ion trap (Normal Scan Rate) using collision-induced dissociation (CID) with normalized collision energy of 35%, activation time of 10ms, activation Q of 0.25, isolation width of 2Th, and automatic gain control target value of 1×10^4^. Ions with charge state 1+ were excluded from precursor selection. Monoisotopic precursor selection was activated. Dynamic exclusion for precursor ions was applied for 45s after 1 fragmentation count and a repeat duration of 30s.

### 4.17 LC-MS/MS data analysis

LC-MS/MS raw data files were used for protein identification using MaxQuant software [85] version 1.5.5.1. Default MaxQuant parameters were used except for the following: protein database was the Mus musculus Uniprot proteome (downloaded on November 19th, 2018); LFQ quantification was selected with default parameters; calculation of iBAQ values was selected; variable modifications, Oxidation(M) and Acetyl(Protein N-term); fixed modifications, Carbamydomethylation(C). The statistical analyses of MaxQuant LFQ intensities were performed by Perseus software [86] version 1.6.14.0, as follows. The dataset was filtered to remove proteins identified as potential contaminants, those only identified by site or by a reverse sequence, and proteins with fewer than two unique peptides. LFQ intensities were Log2-transformed and missing values were replaced using a downshifted normal distribution of the total matrix (width 3, downshift 1.8). The quantitative differences between conditions were evaluated after hierarchical clustering of the data and graphical representation in a volcano plot (Student t-test with false discovery rate of 0.05 and artificial within groups variance S0=1). Protein interaction network and functional classification of the differentially regulated proteins was done using STRING (https://string-db.org) [87].

### 4.17 Statistical analyses

All data is presented as mean ± standard deviation (SD) and were analysed using GraphPad Prism® version 7.04 (GraphPad Software, San Diego, US). All *in vitro* experiments were conducted on 3 separate occasions, and *in vivo* studies had 5 animals per group. The data was tested for normality using the D’Agostino and Pearson omnibus normality test and was analysed using unpaired students t-test, or a Mann-Whitney test for non-parametric distribution. An ANOVA with Dunnett’s multiple comparison post-test or the Kruskal–Wallis test with Dunn’s multiple comparison post-test was used for comparing three or more groups, depending on the normality of the data distribution. All analyses were performed two-tailed with a critical p-value of 0.05.

## Supplementary Materials

Supplementary Figure 1-2

Supplementary Tables 1-3

## Author Contributions

KMS, PO, JTE and AG designed the study. KMS, LT, AH, SCM and AEAM performed the experiments. KMS, AH, LT, SCM, AEAM and AW analysed the data. All authors contributed to interpretation of the analyses, edited and approved the final manuscript.

---

**Supplementary Figure 1**. Historphometric analyses of the tibial endocortical bone (n= 5 animals per group) was performed by measuring A) osteoblast number per bone perimeter (N.Ob/B.Pm), and B) osteoclast number per bone perimeter (N.Oc/B.Pm). Osteoclasts were generated from the mononuclear haematopoietic cell population isolated from long bone marrow of 8-week old female mice and cultured on dentine disks. At the onset of resorption which represents a mature osteoclast phenotype, the cells were treated with vehicle or 10μM A740003 for a further 7 days. At the end of the culture, the cells were fixed, TRAP-stained and quantified for C) total osteoclast number, D) percentage resorbing osteoclasts, E) percentage resorption area, and F) resorption per resorbing osteoclast. G) Representative images of TRAP-stained dentine disks. The resorbing osteoclasts (black star), non-resorbing osteoclasts (yellow star) and resorption pits (arrow) have been highlighted. H) SaOS-2 cells were cultured in osteogenic media for 7 days with vehicle or 10μM A740003. The media was supplemented with inorganic phosphates for the last 2 days of the culture. The cells were then fixed and percentage area of mineralisation assessed using Alizarin red staining. All *in vitro* data are from 3 separate biological repeats. The data has been presented as mean ± SD and analysed using unpaired t-test.

**Supplementary Figure 2**. A) μCT analyses of osteolytic lesions in tibias of 4T1 tumour bearing BALB/c P2RX7^−/-^ mice treated daily with P2RX7 antagonist (10mg/kg A740003) or vehicle controls (n=5 per group); AMC are non-tumour bearing age-matched controls. B) E0771 cells were treated with varying concentrations of BzATP (0-300μM), and conditioned media was collected. Western blotting for LOX shows the effect of P2RX7 activation on LOX secretion. The band for N-glycosylated inactive pro-LOX (gly-LOX) is seen at ∼58kDa and the mature active LOX is 32kDa. C) P2RX7 activation with 300μM BzATP changes the ratio of inactive gly-LOX to mature LOX in the conditioned media, and this is inhibited in presence of A740003. The data has been presented as mean ± SD and analysed using one-way ANOVA. *P<0.05 and ***P<0.0001.

