## Supplemental Table 3 for "P2RX7 inhibition reduces breast cancer induced osteolytic lesions - implications for bone metastasis"

Supplementary Table 3. Protocols for immunohistochemistry

|  | **Protocols** | | | | |
| --- | --- | --- | --- | --- | --- |
| **Antibodies** | Ki67  (Abcam, ab15580) | Caspase-3  (R&D Systems, AF835) | Endomucin  (Santa Cruz, sc-53940) | F4/80  (Bio-Rad,MCA497) | GLUT-1  (Abcam, ab652) |
| **Antigen retrieval** | Tris-EDTA pH 8 at 95°C for 25 min | Citrate Buffer pH 6 at 70°C – 30min | Citrate Buffer pH 6 at 95°C – 25 min | Proteinase-K at 3µg/mL for 20 min | Citrate Buffer pH 6 at 95°C – 25 min |
| **Vectastain ABC Kit** | Goat - anti Rabbit IgG | Goat - anti Rabbit IgG | Rabbit – anti Rat IgG | Rabbit – anti Rat IgG | Goat - anti Rabbit IgG |
| **1° Antibody Concentration** | 3µg/mL | 5µg/mL | 2µg/mL | 20µg/mL | 1:250 dilution of antiserum |
