## Supplementary figures and images for "P2RX7 inhibition reduces breast cancer induced osteolytic lesions - implications for bone metastasis"

### Supplemental Figures

**Supplementary Figure 1**

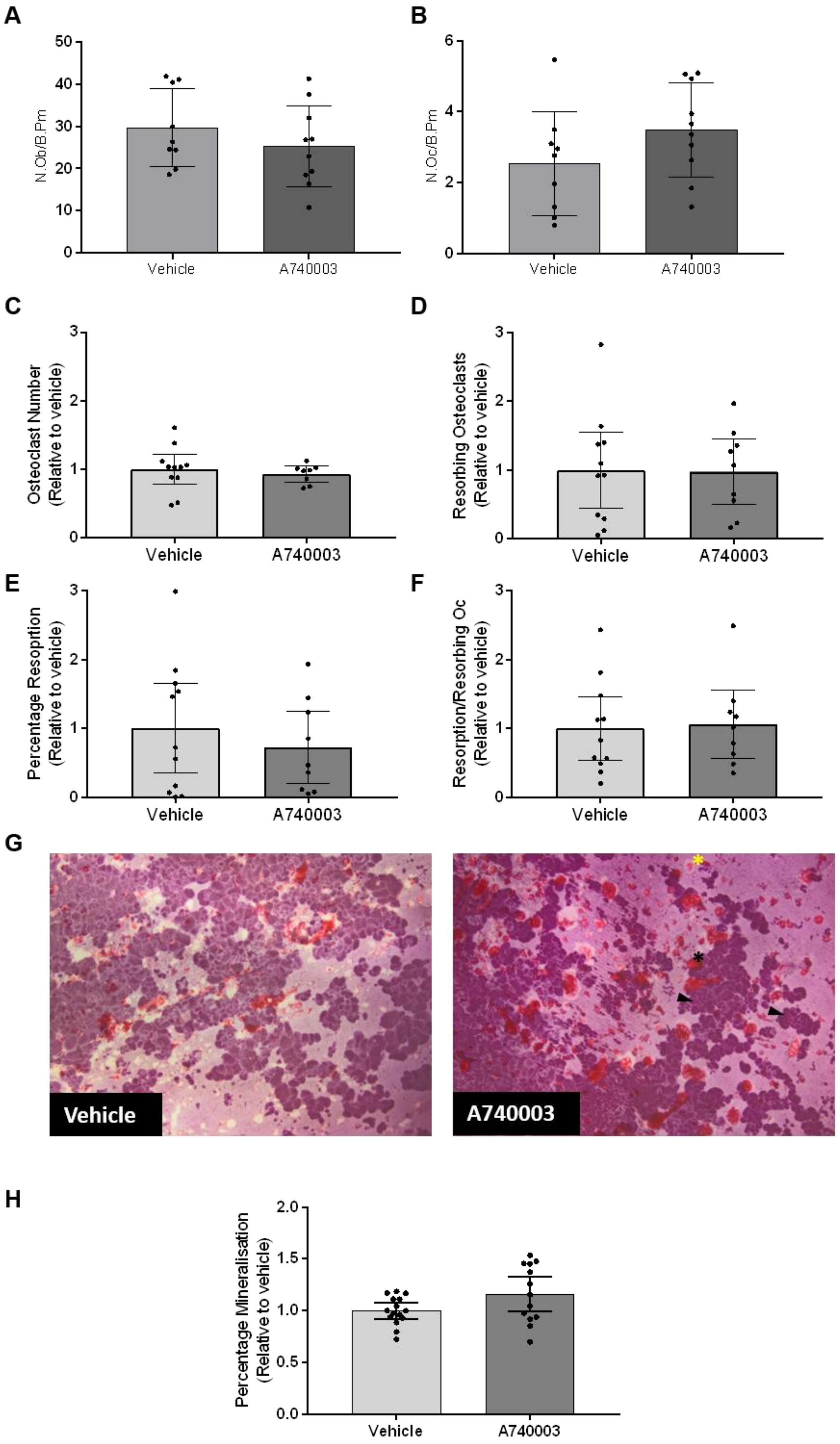

**Supplementary Figure 2**

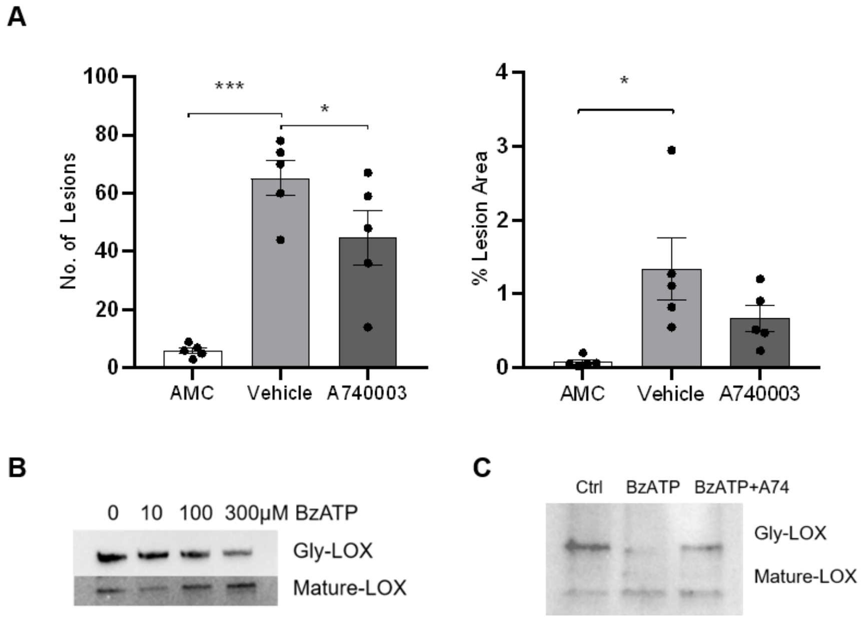
